# The single-cell make-up of adult diffuse glioma based on the 2021 WHO classification

**DOI:** 10.1101/2024.07.23.604812

**Authors:** Iyad Alnahhas, Allison Kayne, Mehak Khan, Wenyin Shi

**Affiliations:** Division of Neuro-Oncology, Department of Neurology, Thomas Jefferson University; Sidney Kimmel Medical College, Thomas Jefferson University; Department of Radiation Oncology, Thomas Jefferson University

## Abstract

**Introduction:** Single-cell RNA sequencing (scRNA-seq) has helped to elucidate the cellular composition of cancer and its microenvironment. Recent scRNA-seq studies have highlighted the heterogeneity of glioblastoma (GBM). Moreover, single-cell GBM analyses have proposed resemblance of GBM cells to radial glia and outer radial glia supporting the hypothesis that remnants of developmental tissue get reactivated in cancer. A recent study isolated neural progenitor cells (NPCs) from developing fetal human brain (gestational week 17–19) and classified NPCs based on their expression of THY1, CD24 and EGFR. Ventricular radial glia are THY1-CD24-EGFR+ whereas outer radial glia are THY1-CD24-EGFR-. Early neuron precursors are CD24+THY1-EGFR+ and glial progenitor cells (GPCs) are THY1+EGFR+. GPCs give rise to THY1+EGFR+PDGFRA+ pre-oligodendrocyte progenitor cells. The importance of EGFR in NPCs again highlights the resemblance to glioma.

**Methods:** We aimed to apply the classification above in IDH mutant astrocytoma and oligodendroglioma as well as IDHwt glioblastoma samples. We used three publicly available datasets: Wang (paired 74 IDHwt primary and recurrent samples), Tirosh (6 primary oligodendroglioma samples) and Venteicher (10 primary IDH mutant astrocytoma).

**Results:** In IDH mutant astrocytoma, 82.63% of cells express THY1+ (mostly EGFR+PDGFRA+) and 10.76% of cells are THY1-CD24-EGFR+. In oligodendroglioma, 75% of cells are THY1+ (mostly EGFR+PDGFRA+) and 12.07% are THY1-CD24-EGFR+. In IDHwt EGFR amplified primary GBM samples, 87.5% of cells are THY1-CD24-EGFR+. This percentage drops to 70.4% in the recurrent setting. THY1-CD24-EGFR-cells increase from 9.7% to 23.1% at recurrence. In IDHwt EGFRwt primary GBM samples, 48.6% of cells are THY1-CD24-EGFR+ and 44.15% are THY1-CD24-EGFR-. In the recurrent setting, 43.26% of cells are THY1-CD24-EGFR+ and 49.58% are THY1-CD24-EGFR-.

**Conclusion:** IDH mutant gliomas and IDHwt glioblastoma express different progenitor cell markers. THY1 is highly expressed in IDH mutant gliomas.

## Introduction

Astrocytomas include IDH1/2 mutant and IDH wild-type (IDHwt) subtypes. Glioblastoma (GBM) is the most common malignant brain tumor and represents IDHwt astrocytoma grade IV (1).

Little advances have been made in the treatment of glioblastoma despite enormous research efforts. In fact, glioblastoma was among the first cancers to be profiled by The Cancer Genome Atlas Project (TCGA) (2). Since then, we have well understood the epigenetics and genetics of glioblastoma at the DNA and transcriptomic levels. Receptor tyrosine kinase (RTK /RAS) pathway alterations (e.g. via amplification of epidermal growth factor receptor (EGFR) are among the most common altered pathways in glioblastoma. Chromosome 7 gain (containing EGFR) and chromosome 10 loss (containing PTEN among other tumor suppressor genes) are thought to be early initiating events in the gliomagenesis of IDHwt GBM (3).

Taking this information into account, and given the prognostic relevance, certain molecular markers have been added as elements for CNS tumor grading based on the 2021 WHO classification (4), whereas grading was traditionally solely based on histological features. For IDHwt astrocytoma specifically, the presence of TERT promoter mutation, EGFR amplification, and/or chromosome 7 gain/10 loss upgrade the tumors to molecular grade IV.

Advanced technologies such as single-cell RNA (scRNA) sequencing /single-nucleus RNA sequencing (snRNA-seq) and spatial transcriptomics have helped to further elucidate the cell-type composition as well as the complexity of this deadly cancer and its microenvironment. Recent scRNA/single-nucleus RNA sequencing (snRNA-seq) studies have demonstrated that GBM cells exhibit a high degree of heterogeneity and plasticity and seamless transitions between cellular states (5, 6). Moreover, interestingly, single-cell GBM analyses have proposed resemblance of GBM cells to radial glia and supporting the ‘‘embryonic rest’’ hypothesis and reactivation of remnants of developmental tissue in cancer (7, 8). A newly published study isolated neural stem and progenitor cells from developing fetal human brain (gestational week (17–19) and classified radial glia and outer radial glia based on their expression of EGFR, CD24 and THY1 (9). Ventricular radial glia (vRG) are CD24-THY1-EGFR+, whereas outer radial glia (oRG) are CD24-THY1-EGFR-. Early neuron precursors are CD24+THY1-EGFR+ and glial progenitor cells (GPCs) are THY1+EGFR+. GPCs are lineage-restricted to astrocytes and oligondedrocytes but not to neurons. THY1+ cells are further classified based on EGFR and PDGFRA expression to pre-OPCs (THY1+EGFR+PDGFRA+) and OPCs (THY1+EGFR-PDGFRA+) that finally give rise to mature oligodendrocytes (THY1+EGFR-PDGFRA-). The importance of EGFR and PDGFRA as a markers for these progenitor cells again highlights the resemblance to markers of gliomagenesis.

Most of the GBM single-cell publications were limited by small sample size, given how expensive these technologies are. Some studies combined IDH mutant and IDHwt samples in the analysis, and some combined primary and recurrent samples. Not all studies reported on the mutations/ copy number change data relevant to the new IDHwt GBM classification. The purpose of this study is to describe the single-cell make-up of GBM based on the WHO 2021 classification. To accomplish this, we use publicly available data from three recent scRNA/snRNA glioma studies: Wang et al (6), Venteicher et al (10) and Tirosh et al (11). We aimed to apply the CD24/THY1/EGFR classification above to IDHwt GBM, IDH mutant astrocytoma and IDH mutant oligodendroglioma, respectively.

## Methods

Data from the Wang, et al study (6) was downloaded using GEO accession GSE174554. The study profiled 86 primary and recurrent GBM specimens with snRNA sequencing. 76 of these samples represent IDHwt GBM. 52 of these samples had patient-matched pair identifiers. The supplementary tables were downloaded from the original paper. Information about EGFR amplification/ mutation status, TERTp mutation as well as chromosome 7 gain/ 10 loss were extracted manually from Figure 1 from the original manuscript into ‘sample_df.csv’ (supplementary material). EGFR and TERTp data were based on the UCSF500 clinical assay panel. Chr7/10 data were derived from snRNA analysis per Wang et al.

**Figure 1.**
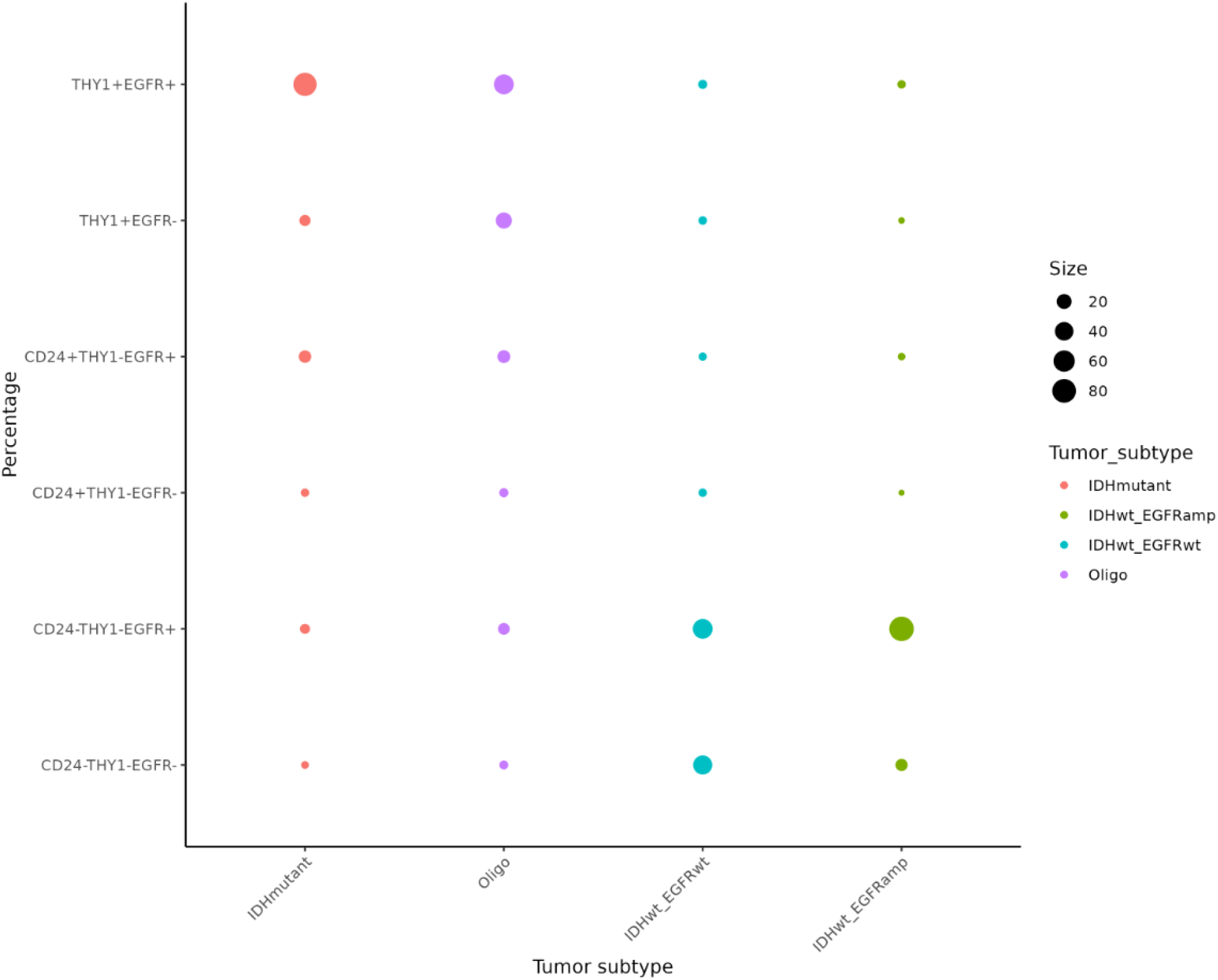
shows the cellular composition IDH mutant astrocytoma, 1p19q co-deleted oligodendrogliomas and IDHwt glioblastoma.

For the Venteicher et al 2017 (10) and Tirosh et al (11) studies, the pre-processed matrix file was downloaded from the 3CA database (12). The 3CA database houses 77 scRNA datasets where the quality control, filtering and cell-type annotation were all consistently applied to all the datasets and made available to download. The Venteicher et al study included 10 IDH mutant astrocytoma samples of various grades: 1 primary grade II, 6 primary grade III, 1 recurrent grade III, 1 primary grade IV and 1 recurrent grade IV. The Tirosh et al study included 6 IDH mutant/ 1p19q co-deleted oligodendroglioma samples: five were grade II and one was “grade II/III”.

We used R 4.3.1 to analyze the scRNA/snRNA datasets. Seurat objects were created for the above studies per the Seurat V5 workflow (13). For the Wang et al study, quality control for was completed per the Seurat workflow. Cells were selected for further analysis after excluding potential empty droplets (cells with less than 200 genes per cells) and doublets or multiplets (cells with more than 2500 genes per cell). Low quality or dying cells were excluded by selecting cells with less than 5% mitochondrial genes. The data was then normalized, highly variable features were selected, the data was scaled and dimensionality reductions were applied. We excluded samples that were comprised of < 10 tumor cells.

For the Wang samples, cell-type annotation (tumor cells versus microenvironment) was extracted from “GSE174554_Tumor_normal_metadata.txt” from GEO. For the Venteicher and Tirosh samples, cell-type annotation was downloaded from 3CA.

## Results

72 of the Wang samples met the quality control criteria above. After quality control, the number of tumor cells ranged from 22-4224 (median 590 cells). Of the 32 primary samples, 9 were EGFR amplified and 23 were EGFR wild-type (EGFRwt), 27 had Chr7 gain/ Chr10 loss while 5 did not, and 16 were positive for the TERTp mutation and 16 were negative. For the Venteicher samples, the number of tumor cells ranged from 112-1273 (median 441 cells). And for the Tirosh samples, the number of tumor cells ranged from 428-1174 (median 638 cells).

### Applying the CD24/THY1/EGFR classification to astrocytoma

In primary EGFR amplified samples, 87.51% of cells are CD24-THY1-EGFR+. The percentage drops to 70.39% in the recurrent setting. THY1-CD24-EGFR-cells increase from 9.76% to 23.11% at recurrence. THY1+ and CD24+ cells comprise <2% of cells each.

In primary EGFRwt samples, 48.63% of cells are CD24-THY1-EGFR+. The percentage drops to 43.27% in the recurrent setting. THY1-CD24-EGFR-cells increase from 44.15% to 49.58% at recurrence. THY1+ cells comprise 4.3% and 3.8% and CD24+ cells comprise 2.92% and 3.35% in the primary and recurrent settings, respectively.

In IDH mutant astrocytoma, on the other hand, 82.64% of cells overall are THY1+, while 12.31% are THY1-CD24+. Unlike IDHwt GBM, only 5.05% of cells are THY1-CD24-. More specifically, and regardless of the tumor grade, most cells were THY+EGFR+PDGFRA+ and comprised 79.05%-82.47% of primary IDH mutant samples. In the one grade III recurrent sample, THY1-CD24-cells increased to 11.07% and in the one grade IV recurrent sample, CD24+ cells increased to 28.54%. Of the THY1+ cells, THY1+EGFR+PDGFRA-cells (precursor to pre-OPCs) increased from an average of 10.27% in primary grade III samples to 13.22% in the recurrent grade III sample, and from 18.26% in the primary grade IV sample to 74.68% in the recurrent grade IV sample. THY1+EGFR+PDGFRA+ cells decreased from an average of 81.64% in the primary grade III samples to 57.93% in the recurrent grade III sample, and from 79.05% in the primary grade IV sample to 10.58% in the recurrent grade IV sample.

Similarly, of 1p19q co-deleted oligodendrogliomas, 75.09% of cells were THY1+, 14.69% of cells were CD24+ and 10.73% of cells were THY1-CD24-. Figure 1 shows the cellular composition of all primary IDH mutant astrocytoma, 1p19q co-deleted oligodendroglioma and IDHwt glioblastoma samples.

### The effect of chromosome 7 gain/ 10 loss on the CD24/THY1/EGFR classification in IDHwt GBM

Of the 9 primary EGFR amplified IDHwt GBM samples in the Wang cohort, 8 had Chr 7 gain/ 10 loss. On the other hand, of the 23 primary EGFRwt GBM samples, 19 had Chr 7 gain/ 10 loss. Therefore, and in order to assess the pure effect of Chr7 gain/ 10 loss on the cellular composition in IDHwt GBM -isolated from the EGFR amplification effect-we compared the cellular composition of the 19 EGFRwt samples with Chr 7 gain/ 10 loss to the 4 EGFR wt samples that were negative for Chr 7 gain/ 10 loss.

Primary Chr7 gain/ 10 loss negative samples are composed of 85.69% of CD24-THY1-EGFR-cells and 10.84% of CD24-THY1-EGFR+ cells. The percentage becomes 61.54% and 33.18% respectively in the recurrent setting. On the other hand, primary Chr7 gain/ 10 loss positive samples include on average 46.5% of CD24-THY1-EGFR-cells and 45.87% of CD24-THY1-EGFR+ cells. The percentage becomes 43.12% and 49.56% respectively in the recurrent setting.

### TERT expression per TERTp mutation status

Of the 26 pairs of samples, TERTp mutaton status only changed once from absent to present in the recurrent setting. Overall, TERT expression was very low regardless of the TERTp mutation status. Only 1% of cells express TERT when TERT promoter mutation is negative and 1.4% of cells express TERT when TERT promoter mutation is positive.

### Find differentially expressed markers between EGFR+ and EGFR-cells

We then used Seurat’s FindMarkers function to find differentially expressed genes between cells expressing EGFR and cells not expressing EGFR. We performed these analyses in EGFR amplified and EGFRwt as well as primary and recurrent samples separately. Supplementary tables 2-5 include data on the differentially expressed genes that met a significant adjusted p-value of <0.01 and log fold-change of the average expression > 1 or < -1 in EGFR amplified primary and recurrent samples, and EGFRwt primary and recurrent samples.

### Functional enrichment analysis of differentially expressed genes

Gene Set Enrichment Analysis (GSEA) of the differentially expressed genes between EGFR+ and EGFR-cells was applied. Within the EGFR amplified samples in the primary setting (supplementary table 6), and as expected, the VERHAAK_GLIOBLASTOMA_CLASSICAL set is highly enriched in EGFR+ cells (normalized enrichment score (NES) -2.3, adjusted p value 0.00016). On the other hand, the VERHAAK_GLIOBLASTOMA_ MESENCHYMAL set is enriched in EGFR-cells (NES 1.97, adjusted p value 0.0008). Seven hypoxia gene sets are enriched in the EGFR-cells but none in EGFR+ cells.

Within the EGFR amplified samples in the recurrent setting (supplementary table 7), the VERHAAK_GLIOBLASTOMA_CLASSICAL set remains enriched in EGFR+ cells (NES - 3.38, adjusted p value 0.0000000001). However, VERHAAK_GLIOBLASTOMA_PRONEURAL becomes enriched in EGFR-cells (NES 3, adjusted p value 7.741667e-08). Intrestingly, eleven hypoxia gene sets are enriched in EGFR+ cells and one hypoxia gene set is enriched in EGFR-cells.

Within the EGFRwt samples in the primary setting (supplementary table 8), the VERHAAK_GLIOBLASTOMA_CLASSICAL set is also enriched in EGFR+ cells (NES – 2.9, adjusted p value 1.499300e-08). VERHAAK_GLIOBLASTOMA_MESENCHYMAL and VERHAAK_GLIOBLASTOMA_PRONEURAL are enriched in EGFR-cells (NES 2.35, adjusted p value 3.466847e-07) and (NES 1.52, adjusted p value 4.842140e-02), respectively. Twenty hypoxia gene sets are enriched in the EGFR-cells but none in EGFR+ cells.

Finally, within the EGFRwt samples in the recurrent setting (supplementary table 9), the VERHAAK_GLIOBLASTOMA_CLASSICAL and VERHAAK_GLIOBLASTOMA_PRONEURAL sets are enriched in EGFR+ cells (NES -2.78, adjusted p value 1.238314e-08) and (NES -2.07, adjusted p value 2.092531e-02), respectively. On the other hand, the VERHAAK_GLIOBLASTOMA_ MESENCHYMAL set is enriched in EGFR-cells (NES 3.03, adjusted p value 1.215676e-08). Twenty hypoxia gene sets are enriched in the EGFR-cells but none in EGFR+ cells.

## Discussion

Advanced technologies such as next-generation sequencing and single-cell RNA (scRNA) sequencing have helped to elucidate the cell-type composition as well as the complexity of cancer and its microenvironment. Moreover, single-cell glioma analyses have proposed resemblance of GBM cells to radial glia and outer radial glia. The purpose of this study is to describe the single-cell make-up of GBM based on the WHO 2021 classification. To this purpose, we used a newly described classification of neural stem and progenitor cells from developing fetal human brain using the markers CD24, THY1 and EGFR. Ventricular radial glia (vRG) are CD24-THY1-EGFR+, whereas outer radial glia (oRG) are CD24-THY1-EGFR-. Early neuron precursors are CD24+THY1-EGFR+ and GPCs are THY1+EGFR+. GPCs are lineage-restricted to astrocytes and oligondedrocytes but not to neurons.

We find that primary IDHwt GBM samples are mainly composed of cells that express markers similar to ventricular and outer radial glia. Chromosome 7 gain and EGFR amplification drive a higher proportion of CD24-THY1-EGFR+ cells. In primary EGFR amplified samples, 87.51% of cells are CD24-THY1-EGFR+. Whereas THY1+ and CD24+ cells comprise <2% of cells. EGFRwt GBM that lacks chromosome 7 gain is composed of 85.69% of CD24-THY1-EGFR-cells, on the opposite extreme. The proportion of THY1+ and CD24+ cells are higher (4.3% and 3.8%, respectively) in EGFRwt GBM. The remaining cells are roughtly evenly split between CD24-THY1-EGFR+ and CD24-THY1-EGFR-cells. The difference is striking in IDH mutant astrocytoma and oligodendroglioma. In IDH mutant astrocytoma 82.64% of cells overall are THY1+ and 12.31% are THY1-CD24+. Similarly, of 1p19q co-deleted oligodendrogliomas, 75.09% of cells were THY1+, 14.69% of cells were CD24+. Most of the THY1+ cells in IDH mutant glioma are also EGFR+PDGFRA+, an expression profile similar to pre-OPCs. This data suggests that IDH mutant gliomas and IDHwt glioblastoma express different progenitor cell markers and may have a different cell of origin. THY1 is highly expressed in IDH mutant gliomas.

Another interesting phenomenon is that, consistently, a different cell population than the dominant one in the primary setting becomes more prevalent in the recurrent setting. CD24-THY1-EGFR-cells increase from 9.76% to 23.11% at recurrence in EGFR amplified GBM, and from 44.15% to 49.58% at recurrence in EGFRwt GBM. CD24+ and THY1+EGFR+PDGFRA-increase in proportion in recurrent IDH mutant astrocytoma.

Gene set enrichment analysis shows that ther VERHAAK_GLIOBLASTOMA_CLASSICAL set is consistently enriched in EGFR+ cells regardless of EGFR amplification status. The VERHAAK_GLIOBLASTOMA_ MESENCHYMAL set is enriched in EGFR-cells in EGFRwt primary and recurrent samples. More hypoxia gene sets are enriched in EGFR-cells in EGFRwt primary and recurrent samples compared to EGFR amplified samples.

Finally, TERT expression was very low, even in the presence of TERTp mutation (1.4% of cells). This is consistent with recent literature that suggests that unlike the *TERT* promoter mutation, which was seen throughout the entire tumor, *TERT* expression was detected only in a subset of cells (14).

## Supporting information

Suppelmentary table 1

sessionInfo

Suppelmentary table 4

Suppelmentary table 5

Suppelmentary table 2

Suppelmentary table 3

Suppelmentary table 8

Suppelmentary table 9

Suppelmentary table 6

Suppelmentary table 7

## Supplementary material

Suppelmentary table 1: Wang et al ‘sample_df.csv’

Suppelmentary table 2: Differentially expressed genes between EGFR+ and EGFR-cells in EGFRamp primary samples.

Suppelmentary table 3: Differentially expressed genes between EGFR+ and EGFR-cells in EGFRamp recurrent samples.

Suppelmentary table 4: Differentially expressed genes between EGFR+ and EGFR-cells in EGFRwt primary samples.

Suppelmentary table 5: Differentially expressed genes between EGFR+ and EGFR-cells in EGFRwt recurrent samples.

Suppelmentary table 6: Gene Set Enrichment Analysis (GSEA) of the differentially expressed genes between EGFR+ and EGFR-cells in EGFRamp primary samples.

Suppelmentary table 7: Gene Set Enrichment Analysis (GSEA) of the differentially expressed genes between EGFR+ and EGFR-cells in EGFRamp recurrent samples.

Suppelmentary table 8: Gene Set Enrichment Analysis (GSEA) of the differentially expressed genes between EGFR+ and EGFR-cells in EGFRwt primary samples.

Suppelmentary table 9: Gene Set Enrichment Analysis (GSEA) of the differentially expressed genes between EGFR+ and EGFR-cells in EGFRwt recurrent samples.

<u>sessionInfo:</u>

The output from running ‘sessionInfo’ details all the R packages and versions used in this script

## Code availability

All code used is available on GitHub (https://github.com/iyadalnahhas/scRNA_astrocytoma)

## Notes

### Competing Interest Statement

The authors have declared no competing interest.

